# The Isolation and Identification of Fungi Gathered from Districts in Bangkok, Thailand Where Dengue Fever Is at Epidemic Levels

**DOI:** 10.1101/518241

**Authors:** Ladawan Wasinpiyamongkol, Panan Kanchanaphum

## Abstract

**Background:** The *Aedes* mosquito is a major vector of many important diseases such as dengue, chikungunya, Zika, and yellow fever. Biological methods of controlling mosquitos are desirable because they are ecologically friendlier, safer, and more cost effective than chemical and physical methods of controlling mosquitos.

**Methods:** Water samples in the mosquitoes’ breeding containers from districts in Bangkok were collected from the mosquitoes breeding containers situated in seven districts of Bangkok, Thailand. The DNA was extracted from each sample of the isolated fungi. Purified DNA specimens were amplified in a PCR reaction with universal primers of ITS1 and ITS4. All the PCR product was sequencing, alignment and comparing the homologous sequence in GenBank database.

**Results:** Fourteen strains of fungi were isolated. The most commonly found strain was *Penicillium citrinum*, which was discovered in six of the 30 isolated fungi samples.

**Conclusion:** Biological control strategies for the mosquito population should be further investigated because they are considered to be ecologically friendlier, safer, and more cost effective than chemical insecticides.

## Introduction

Mosquitoes are an important vector species for arboviruses that belong to the following three families: Flaviviridae, which causes dengue fever, Bunyaviridae, which causes chikungunya, and Zika Togaviridae, which causes arthritis, encephalitis, and rubella [1, 2]. The *Anopheles* species of mosquitoes, which belongs to the *Aedes* and *Culex* genera, is responsible for the majority of arbovirus transmission. *Aedes aegypti* is a highly anthropophilic species known for transmitting several emerging arboviruses such as dengue, Zika, chikungunya and yellow fever, which have had a significant impact on human public health [3].

Dengue fever outbreaks occur intermittently in Thailand. A reduction in dengue transmission could be achieved by controlling the population density of *Aedes aegypti* and ensuring it is maintained at a level below the critical threshold that results in an epidemic [4]. A variety of chemical, physical, and biological methods have been used to decrease the incidence of vector-borne diseases such as dengue fever transmitted by mosquitoes. However, the use of chemical insecticides has caused resistance in mosquitoes, serious health hazards, and harmful effects on beneficial non-target animals [5]. Biological control methods are ecologically friendlier, safer, and more cost effective than chemical and physical methods, which are more expensive and time-consuming for the regular entomological observation of mosquito breeding sites [3]. Biological control depends on the use of predators such as fish, the larvae of *Toxorhynchites*, and copepods or parasitic organisms such as *Bacillus thuringiensis israelensis* and *Lysinibacillus sphaericus* as well as entomopathogenic fungus targeting disease vectors [3].

The purpose of this research is to discover fungi with entomopathogenic properties that could be used to control the mosquito population. To achieve this objective, the water samples taken from the mosquito breeding containers situated in the dengue-endemic areas in Bangkok, where is the one of the provinces in the list of the highest outbreak area.

## Materials and methods

### Water sample collection

Water samples were collected from the mosquitoes breeding containers situated in seven districts of Bangkok, Thailand where dengue fever was reported to be at endemic levels (Bang Khen, Lat Krabang, Min Buri, Phra Khanong, Rat Burana, Taling Chan, and Thung Khru). The water samples were kept in 250 ml screw capped sterilized bottles that were sealed and then stored at 4°C prior to the isolation procedure.

### Isolation of filamentous fungi

Initially, the water samples were filtered through a 47 mm diameter sterile 0.45 μm membrane cellulose nitrate filter (Whatman, 7141-104; Whatman International Ltd, UK) using Millipore vacuum apparatus in a laminated flow chamber. Then, using sterile forceps, the filter was transferred to a petri dish containing 4 ml of sterile water where it was thoroughly washed for 30 sec. Next, 1 ml of sterile water was cultured on Potato Dextrose Agar (PDA, Difco, BBL / USA) supplemented with Choloramphenicol 50 mg/l and Gentamycine 25 mg/l. Three replicates were used for each water sample. After that, the agar plates were incubated at 28 ± 2°C for 7-10 days and then monitored daily for the appearance of fungal colonies. Finally, the fungal isolates were subcultured separately to obtain pure cultures on PDA for identification.

### Identification of filamentous fungi

Colony descriptions were based on PDA observations under ambient daylight conditions. Microscopic observations and measurements were made from preparations that were mounted in lactic acid. The morphological characteristics of the fungi were septation of hypha, formation, morphology, the branching frequency of fruiting-bodies and conidiophores. Characteristics of the colony form, structure, size, and color were also observed and recorded. Filamentous fungi were identified at the generic level according to morphological characteristics as described in fungal atlases [6, 7].

### The sequencing of fungal strains by ITS

The DNA was extracted from each sample of the isolated fungi by a fungal DNA extraction kit using the manufacturer’s instructions (PrestoTM Mini gDNA Yeast kit; Geneaid, New Taipei City, Taiwan). Purified DNA specimens were amplified in a PCR reaction with universal primers of ITS1: 5’-TCCGTAGGTGAACCTGCGC-3’ and ITS4: 5’-TCCTCCGCTTATTGATATGC-3’. The PCR reaction included 0.4 μM of ITS1 and ITS4 primers, 0.2 mM dNTPs, 1.5 mM MgCl_2_, 1xPCR buffer (50 mM KCl, 10 mM Tris-HCl), 1.25 units of *Taq* DNA polymerase (New England Biolabs). The reaction was carried out in a BIO-RAD MJ Mini Personal Thermal Cycler. The cycle conditions consisted of a single initial denaturation at 94°C for 5 min followed by 35 cycles at 94°C for 30 sec, 50°C for 30 sec, 72°C for 1 min, and a final extension step at 72°C for 5 min. The PCR product size was about 550 bp. All the PCR product was sent to Solution of Genetic Technology, South Korea for sequencing.

The resulting sequences were checked and aligned using the BioEdit 7.0 sequence alignment editor (Isis Pharmaceuticals, Inc., Carlsbad, CA, USA). The sequences were compared with a homologous sequence stored in the GenBank database then evaluated using the BLAST program on the National Center for Biotechnology Information (NCBI) website.

## Results

Thirty fungal isolates were isolated from water samples gathered from seven areas around Bangkok. The results of the rDNA-ITS sequencing show that 550 bp of PCR product was sequenced from all the isolate strains. The resulting sequences were compared with the 18S rDNA BLAST sequence stored at the GenBank database using the BLAST tool. The 14 isolated fungal strains were shown in Fig. 1.

**Fig. 1.**
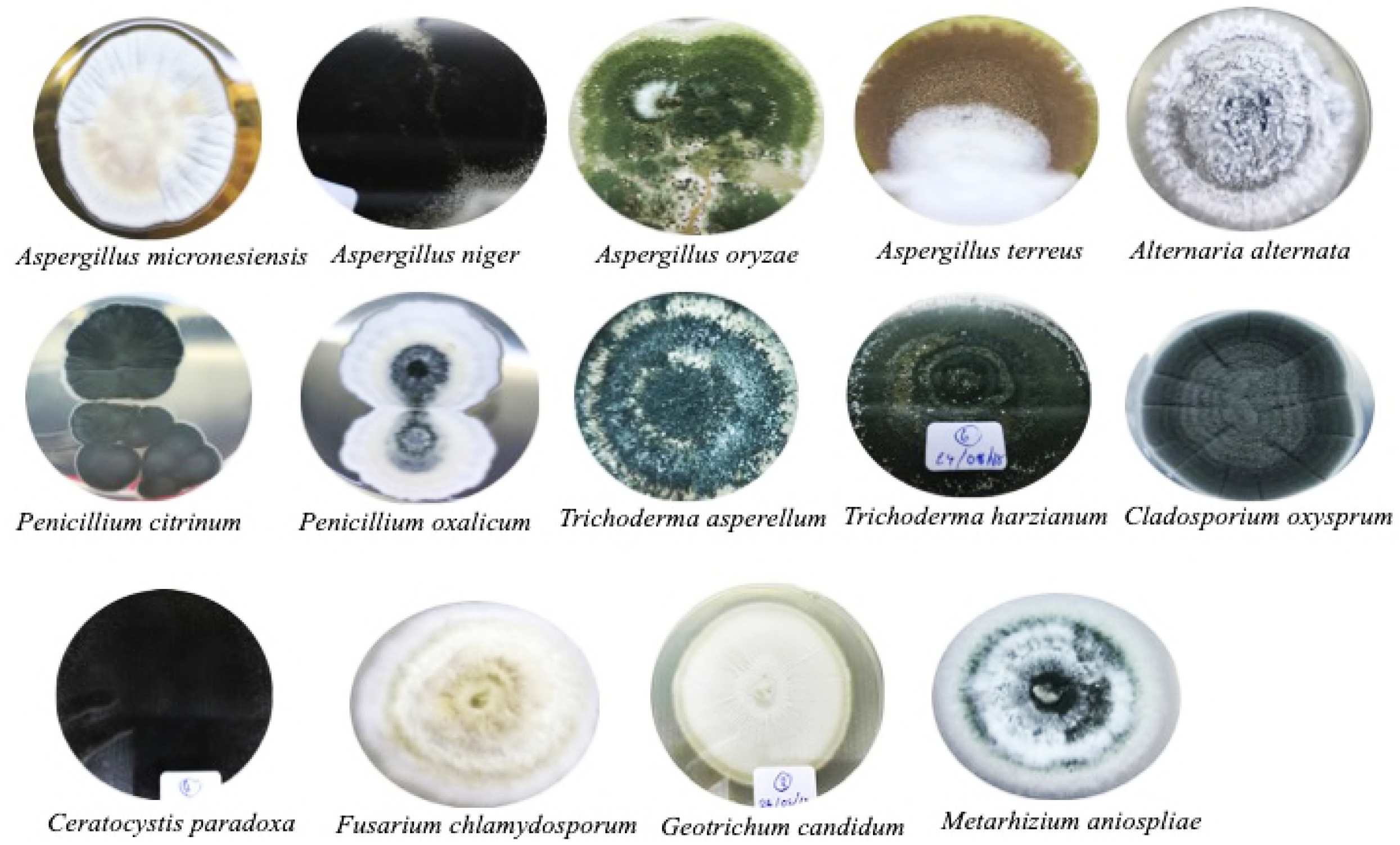
Colony and conidia of isolated fungi on PDA agar medium

*Penicillium citrinum* was found to be the most dominant of all strains isolated. Six isolates of *Penicillium citrinum* were found, accounting for 19.98 % of the total population, as shown in Fig. 2. *Aspergillus oryzae, Fusarium chlamydosporum, Geotrichum candidum, Ceratocystis paradoxa*, and *Trichoderma harzianum* were present in 3.33 % of the total population. *Aspergillus terrrus, Aspergillus niger, Penicillium oxalicum, Cladosporium oxysporum*, and *Metarhizium anisopliae* were present in 6.67 % of the total population. And, *Aspergillus micronesiensis, Alternaria alterata*, and *Trichoderma asperellum* were present in 9.99 % of the total population.

**Fig. 2.**
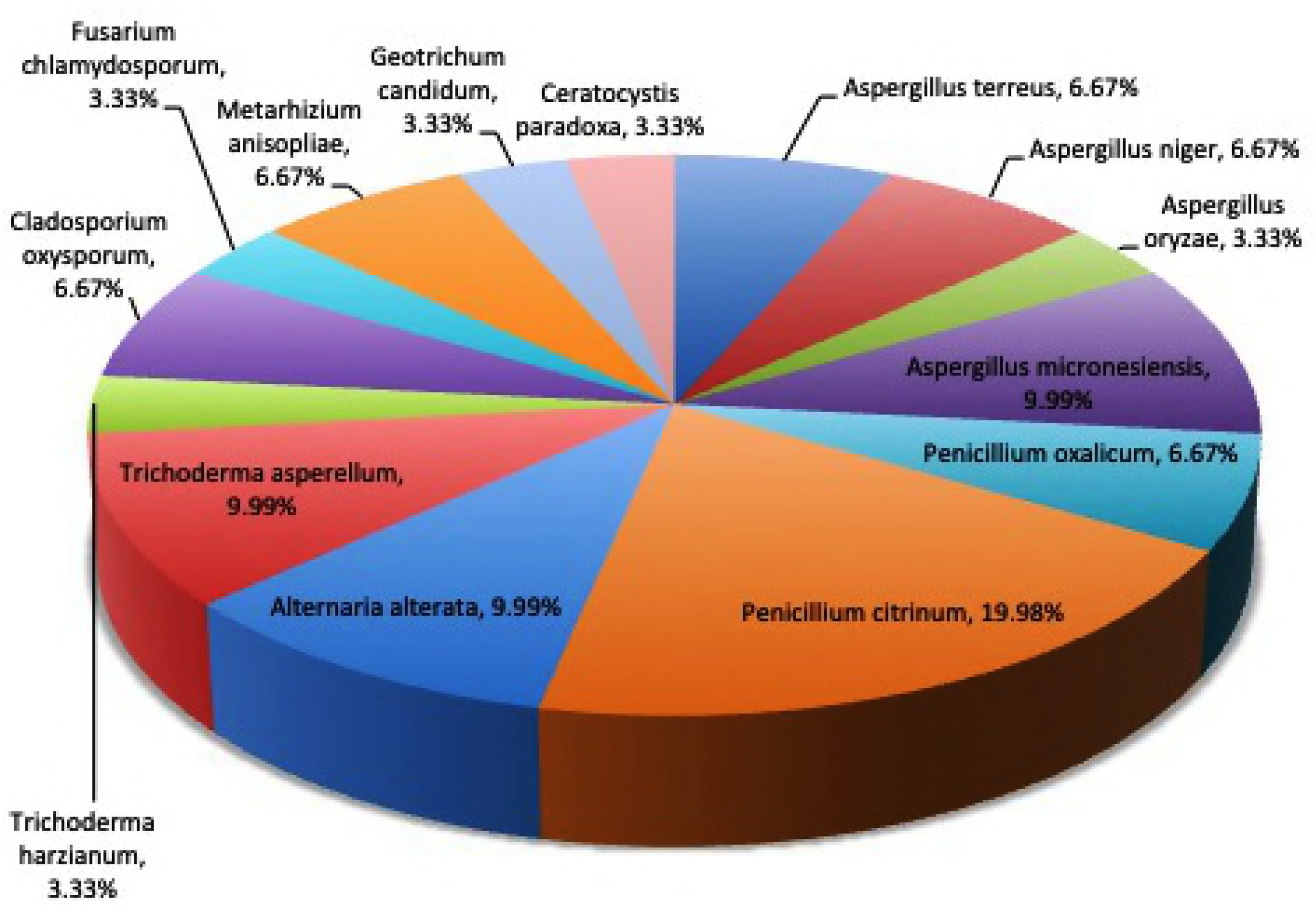
The percentage of isolated strains of fungi gathered from the water samples

## Discussion

Monitoring the diversity of filamentous fungi as part of mosquito vector management could considerably benefit public health. This is the first study in Thailand to isolate and identify fungi gathered from mosquito breeding containers in districts where dengue fever is prevalent around Bangkok. The results show that two strains of the entomopathogenic fungi *Metarhizium anisopliae* and *Penicillium citrinum* could be potential candidates for the biological control of mosquitoes.

Both M. *anisopliae* and *Entomophthora anisopliae* are the fungi that are found in nature throughout the world, which causes disease in various insects by acting as a parasitoid [8]. *M. anisopliae* can infect a wide range of mosquitoes in the genera of *Aedes* and *Culex* [9, 10]. A histological study has reported that the ingestion of conidia eventually leads to blockage of the anatomic structure [11]. The gut of larvae was found to contain high concentrations of conidia, which suggests that it is usually ingested [11]. The mortality of larvae has been related to stress-induced apoptosis, as confirmed by experimental evidence, which demonstrated that treatment with protease inhibitors substantially increased the survival of larvae [12]. Blastospores produced by the *Metarhizium* species have potential for field applications because they exhibit a high degree of host specificity and kill rapidly after penetration [3].

Besides *M. anisopliae*, the other major fungi isolated, which has the potential to kill mosquitoes biologically, was *Penicilium citrinum. P. citrinum* has been found to cause mortality in *Culex quinquefasciatus* or the southern house mosquitoes [13] that acts as a vector of the West nine and Japanese encephalitis viruses [14]. The results of this research show that six of thirty isolated fungi were found to be of the *P. citrinum* species. Therefore, further research should evaluate whether a combination of *M. anisopliae* and *P. citrinum* could be enhanced the effectiveness of the killing mosquito larvae.

Normally, entomopathogenic fungi, endotoxin of *Bacillus thuringiensis israelensis* (*Bti*) and *Lysinibacillus sphaericus* (*Ls*) are used to control the population of mosquitoes. The toxins produced by *Bti* and *Ls* have the advantage of being highly effective as larvicides for various vector species of arboviruses. *Bti* is broadly effective against mosquitoes in the genera of *Aedes, Culex*, and *Anopheles*, whereas the toxicity of *Ls* is limited to *Culex* and *Anopheles* [15].

Research into the application of microorganism insecticides has found that the mosquito-fish, *Gambusia affinis* was compatible with the simultaneous use of other chemical or biological control tools [16]. An experiment in the rice fields of California that mainly focused on *Culex tarsalis* evaluated the release of *G. affinis* followed by treatment with *Bacillus thuringiensis israelensis* [17].

## Conclusions

Entomopathogenic fungi could be used as the potential biological control strategies for mosquito population. Because of these potential fungal isolates are considered to be ecologically friendlier, safer, and more cost effective than chemical insecticides. Further research should be investigated their extracellular metabolites that may be used in integrated the management programs for mosquito population.

## Acknowledgements

We would like to sincerely thank Mr. Stewart Miller for critical correcting English grammar. This work was funded by a grant from Research Institute of Rangsit University, Thailand (Grant no. 63/2560).

